# Deep Learning of High-throughput Transcription Factor–DNA Binding Affinity Data: Quantitative Comparison with Pairwise-Additive Models

**DOI:** 10.64898/2026.05.18.725888

**Authors:** Ke Shen, Zhi Wang, Xiaoliang Sunney Xie

**Affiliations:** Changping Laboratory, Beijing, P. R. China; Biomedical Pioneering Innovation Center (BIOPIC), School of Life Sciences, Peking University, Beijing, P. R. China

**Author notes:** These authors contributed equally to this work (Ke Shen, Zhi Wang).

## Abstract

Transcription factors (TFs) regulate gene expression by binding to specific DNA sequences. Widely used models of TF–DNA binding, such as position weight matrices (PWMs) and position-specific affinity matrices (PSAMs), assume binding free energy is the sum of independent base contributions. However, there is ample evidence that non-additive effects significantly influence TF binding. Here, we utilize data from a high-throughput *in vitro* assay (*ivt*FOODIE) to generate genome-scale TF–DNA dissociation constants (*K*_d_) and systematically evaluate sequence-to-affinity models of increasing complexity. We demonstrate that pairwise additive models exhibit systematic deviations from the measured affinity landscapes. Models incorporating adjacent dinucleotide interactions and deep learning architectures achieve markedly improved agreement with experimental *K*_d_ values. The magnitude of this non-pairwise-additivity depends strongly on the TF family. *In silico* mutation screening reveals widespread, TF-specific long-range interposition dependencies, highlighting the role of energetic coupling across distant positions in target recognition. These results provide a quantitative framework for comparing non-pairwise-additive energetic effects across diverse TFs.

## INTRODUCTION

Transcription factors (TFs) regulate gene expression by recognizing specific DNA sequences with varying strengths of affinity, typically quantified by the dissociation constant, *K*_d_.^1^ A common assumption in modeling TF–DNA recognition is pairwise additivity, which posits that the total binding free energy is the sum of independent energetic contributions from individual bases at each position.^2^ This principle underlies widely used frameworks, including position weight matrices (PWMs)^3^ and position-specific affinity matrices (PSAMs)^4^. These pairwise additive models remain the standard for representing TF specificity due to their interpretability and computational efficiency.^5,6,7^

However, the pairwise additive approximation often fails to capture the intricate physical determinants of TF–DNA binding. Evidence suggests that interdependent effects between nucleotides, where the contribution of one base depends on its neighbors, can significantly modulate affinity.^8,9^ While some studies argue that pairwise additive models provide a reasonable approximation of specificity^10^, the extent to which this assumption holds across diverse TF families and expansive sequence spaces remains a subject of debate. The fundamental challenge lies in determining whether these non-pairwise-additive effects are marginal corrections or essential components of the binding energy landscape.

To move beyond the pairwise additive baseline, various modeling strategies have been developed, ranging from adjacent dinucleotide models^10^ to complex deep learning architectures. In particular, convolutional neural networks (CNNs)^11,12^ offer a powerful, non-parametric framework for capturing high-order sequence dependencies without prior assumptions of positional independence. By learning local sequence features, CNNs can capture the non-linear sequence determinants of binding, providing a complete representation of the sequence–affinity landscape.^13,14,15^

The rigorous evaluation of these models has long been constrained by the nature of available high-throughput data. While technologies such as PBM^16^, HT-SELEX^17^, and ChIP-seq^18^ have yielded vast catalogs of TF binding preferences, they primarily measure relative enrichment or occupancy rather than absolute thermodynamic parameters. This distinction is crucial: without absolute thermodynamic data, it is impossible to strictly quantify the energetic coupling between nucleotides at a physical level. While methods like MITOMI^19^ and HiTS-FLIP^20^ enable high-throughput *K*_d_ measurements, their dependence on specialized instrumentation restricts their applicability and has impeded large-scale mapping of affinity landscapes across TF families.

To bridge this gap, we leverage our recently developed high-throughput assay, *ivt*FOODIE^21^, which provides precise *K*_d_ measurements for 45 TFs across genome-wide DNA libraries. By integrating these absolute thermodynamic data with sequence-to-affinity models of increasing complexity, we systematically map the energetic limits of the pairwise-additive assumption. This framework allows us to evaluate model performance and to quantify the physical magnitude of non-additivity across different TF families. The results provide a direct, genome-scale assessment of the biophysical principles governing TF–DNA recognition, revealing the widespread prevalence of interposition dependencies.

## METHODS

### *ivt*FOODIE Assay and *K*_d_ Estimation

TF–DNA dissociation constants were measured using the high-throughput *ivt*FOODIE assay. Briefly, purified TFs were incubated with a genomic DNA library (derived from ATAC-seq^22^) until equilibrium. The double-stranded cytosine deaminase MGYPDa829 was then introduced to catalyze cytosine-to-uracil (C-to-U) conversion at solvent-exposed sites. Because TF binding sterically protects DNA from deamination, the C-to-U conversion frequency at each binding site inversely correlates with TF occupancy. By measuring these frequencies across a gradient of TF concentrations via high-throughput sequencing, we determined the binding fraction at each concentration. *K*_d_ values were subsequently estimated by fitting these data to the Hill equation (Hill coefficient *n* = 1).^23^

To align model outputs with physical energetics, measured *K*_d_ values were converted to binding affinities defined as –log_10_*K*_d_. Given the thermodynamic identity Δ*G*°=*RT*ln(*K*_d_), this scale ensures that the target metric is linearly proportional to the standard Gibbs free energy of binding. All *K*_d_ values were standardized to molar units prior to conversion.

### Sequence Representation, Model Training, and Evaluation

To systematically evaluate non-additivity in TF– DNA recognition, we implemented four sequence-to-affinity models with increasing representational capacity:

1. PWM: A motif-based baseline generated with STREME^24^ using DNA sequences from *de novo* TF footprints identified in the *ivt*FOODIE data, representing statistically enriched base patterns.
2. PSAM: A linear regression model that assumes independent energetic contributions from individual bases, serving as the pairwise-additive limit.
3. Adjacent Dinucleotide Model: An extension of PSAM incorporating nearest-neighbor coupling terms to capture local non-additivity.
4. CNN: A non-linear architecture comprising two 1D convolutional layers (each followed by batch normalization and ReLU activation) and two fully connected layers.

All models, except PWM, were trained on one-hot encoded DNA sequences to predict –log_10_*K*_d_. Predictive accuracy was assessed using the Pearson correlation coefficient (*r*) between experimental and predicted affinities on held-out DNA sequences.

### *In Silico* Mutations Analysis

To quantify energetic coupling between positions *i* and *j*, we defined the interposition dependency score,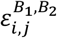, using an *in-silico* double-mutant approach:

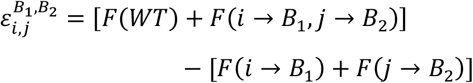

where *F* denotes the CNN-predicted affinity (– log_10_*K*_d_), WT is the consensus binding sequence, and *B* ∈ {*A, C, G, T*} is the substituted base.

To ensure the robustness of *ε* across various sequence contexts, for each TF, we embedded the mutated core motifs into 256 distinct flanking sequences that exhibited the highest experimental affinities. For each position pair (*i, j*), we calculated the signed mean *ε* across all flanking contexts and possible base substitutions. A non-zero *ε* indicates a deviation from the pairwise additive assumption, reflecting energetic epistasis.

## RESULTS

### Non-Pairwise-Additive Models Improve TF–DNA Affinity Prediction

To systematically evaluate the pairwise additive approximation in TF–DNA recognition, we benchmarked four sequence-to-affinity models of increasing architectural complexity against genome-scale *K*_d_ measurements from the *ivt*FOODIE assay (Methods). Our dataset encompasses absolute binding affinities for 45 diverse TFs. As conceptually outlined in Figure 1, the PWM and PSAM served as our pairwise additive baselines. To capture non-pairwise-additive sequence dependencies, we extended the PSAM framework to include adjacent dinucleotide interactions. Concurrently, we deployed a CNN to map higher-order, non-linear relationships. All predictive performances were rigorously evaluated on held-out test sequences to ensure robust generalization (Methods).

**Figure 1.**
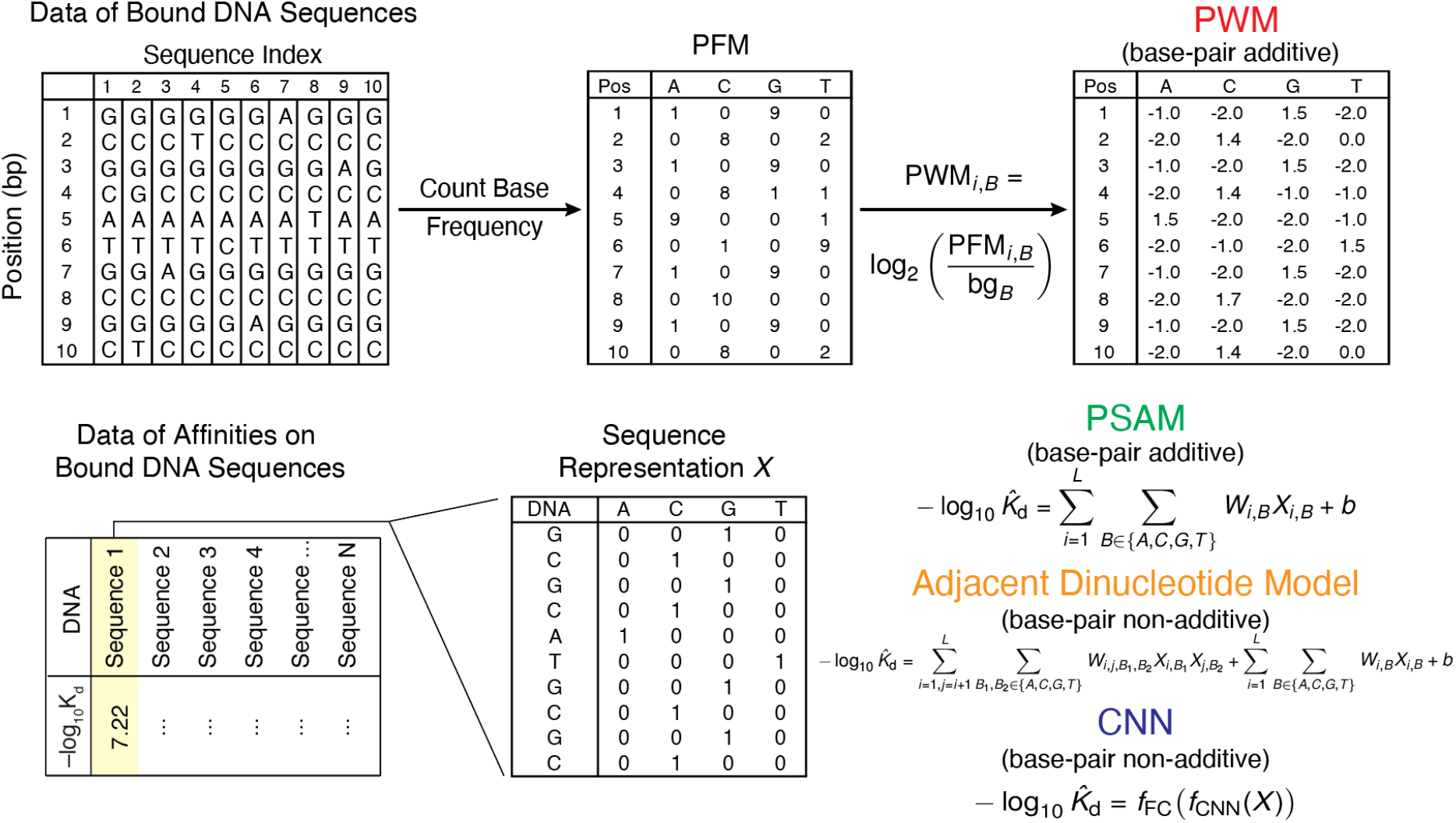
Workflows for modeling TF–DNA binding using PWM, PSAM, adjacent dinucleotide model, and CNN approaches. Top: Generation of a PWM using bound DNA-sequence data. The aligned input sequences are first summarized into a position frequency matrix (PFM), which is then converted into a PWM by calculating the base-2 log-likelihood ratio of the observed frequencies relative to the background probabilities (bg_*B*_). Here, *i* denotes the position within the aligned DNA sequences, and *B* represents the specific nucleotide base (A, C, G, or T). Bottom: Training pipelines for predictive models, utilizing affinity (–log_10_*K*_d_) data on bound DNA sequences. The input DNA sequences are mapped into numerical representations (*X*) via one-hot encoding. These matrices are fed into distinct models to predict affinity: the PSAM model approach relies on linear regression (assuming additive base-pair contributions), whereas the CNN model captures non-pairwise-additive energetic coupling. In the PSAM equation, − log_10_ 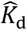 represents the predicted affinity, *W*_*i,B*_ is the weight matrix (which forms the resulting PSAM), and *b* is the bias term. In the adjacent dinucleotide model, the predicted affinity is computed via linear and adjacent non-linear terms. In the CNN model, the predicted affinity is computed via nested non-linear functions, where *f*_CNN_ and *f*_FC_ denote the learned transformations of the convolutional and fully connected layers, respectively.

We first examined representative TFs (SP1, YY1, and CTCF) to visualize the impact of model complexity on predictive accuracy (Figure 2). The traditional PWM exhibited the weakest correspondence with experimental affinities, characterized by broad scatter and low correlation coefficients. By directly regressing on –log_10_*K*_d_, the PSAM model substantially tightened the prediction scatter, capturing a greater portion of the affinity landscape. For each representative TF, incorporating adjacent dinucleotide interactions further enhanced accuracy, while the non-linear CNN visually achieved the strongest agreement with experiment.

**Figure 2.**
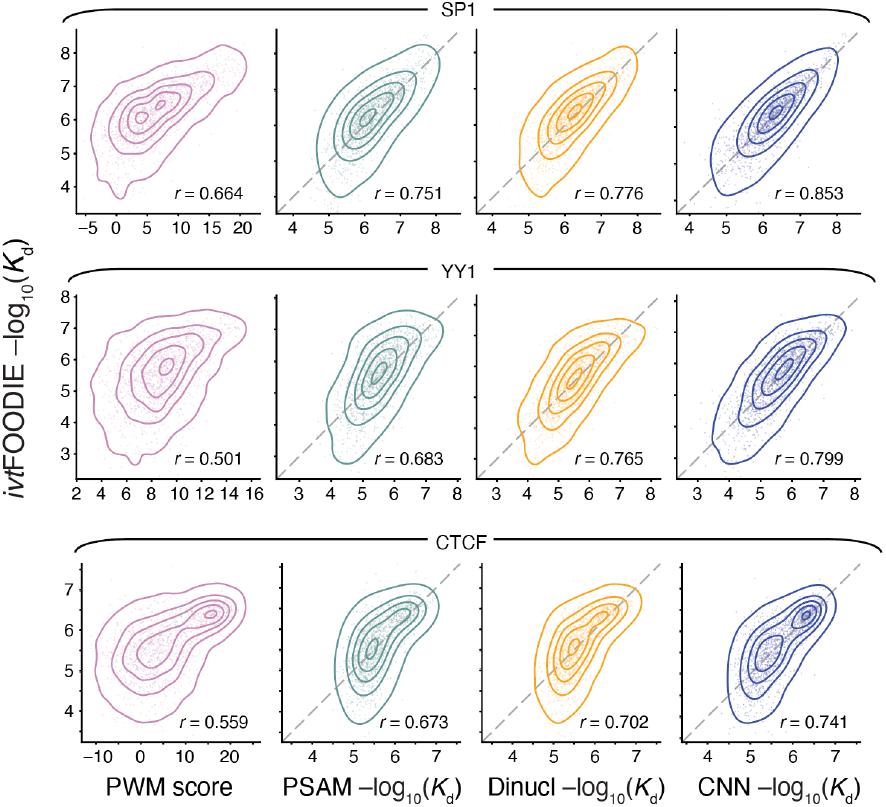
Comparison of model predictions with experimental binding affinities for representative TFs. Scatter plots illustrate the correlation between experimental affinity (–log_10_*K*_d_) and the predicted scores (x-axis) for three distinct models: PWM (left), PSAM (middle left), adjacent dinucleotide model (middle right) and CNN (right), using SP1, YY1, and CTCF as representative examples. Pearson correlation coefficients (*r*) are shown in each panel. For PSAM, adjacent dinucleotide, and CNN models, gray dashed lines indicate the diagonal line. The progressive tightening of the scatter distributions from PWM to CNN (left to right) indicates improved prediction accuracy as increasingly complex sequence dependencies are incorporated.

This qualitative observation is supported by global statistics across the full 45 TF dataset s (Figure 3). The pairwise additive baselines, PWM and PSAM, achieved median Pearson’s *r* of 0.513 and 0.641, respectively. Relaxing the pairwise additive constraint significantly improved predictive power, with the adjacent dinucleotide model increasing the median *r* to 0.701. The CNN achieved the highest global performance with a median *r* of 0.751. Thus, while pairwise additive contributions establish the core binding motif, incorporating nearest-neighbor and high-order sequence dependencies provides a more faithful mathematical representation of the TF–DNA energetic landscape.

**Figure 3.**
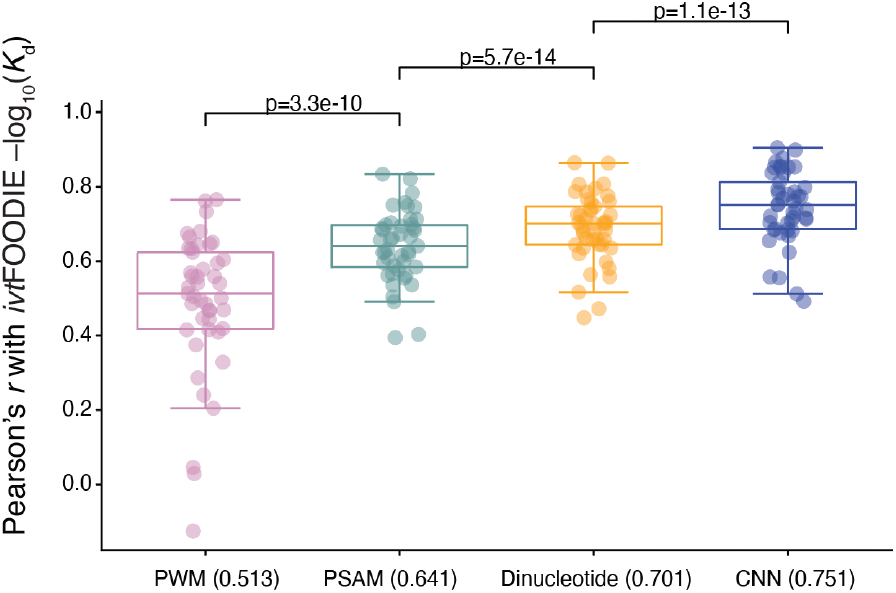
Non-pairwise-additive models significantly outperform pairwise additive models in predicting TF– DNA binding affinities. Boxplots display the performance distributions of the PWM, PSAM, adjacent dinucleotide and CNN models across all evaluated TFs. Performance is quantified by Pearson’s *r* between the experimental –log_10_*K*_d_ values and the respective model predictions. Each dot represents the prediction accuracy for an individual TF, with the median *r* value indicated in parentheses in the x-axis ticks. P values labeled above the boxplot indicate statistical significance calculated by the Wilcoxon signed-rank test.

### The Extent of Non-Pairwise-Additivity Varies across TF Families

TFs recognize DNA sequences through diverse DNA-binding domain (DBD) architectures. We hypothesized that the magnitude of non-pairwise-additivity is TF-family-dependent. To quantify this, we used the performance gap (Δ*r*) between the CNN and PSAM models as a macroscopic metric for non-pairwise-additive energetic effects (Figure 4). We observed striking family-specific divergence. For instance, HOX family homeodomains (e.g., HOXA1, HOXA6, HOXB5) exhibited only marginal non-pairwise-additivity, tightly adhering to the pairwise additive approximation. In contrast, AP-2 family members (TFAP2B, TFAP2C) displayed large predictive gains (Δ*r* ≈ 0.3) with the CNN, indicating profound non-pairwise-additive coupling. The C2H2 zinc finger family, the largest in our dataset, generally exhibited moderate non-pairwise-additivity (Δ*r* ≈ 0.1). Thus, the energetic binding constraints, expressed here as non-pairwise-additive interaction effects are intrinsically linked to the physical structure of the TF.

**Figure 4.**
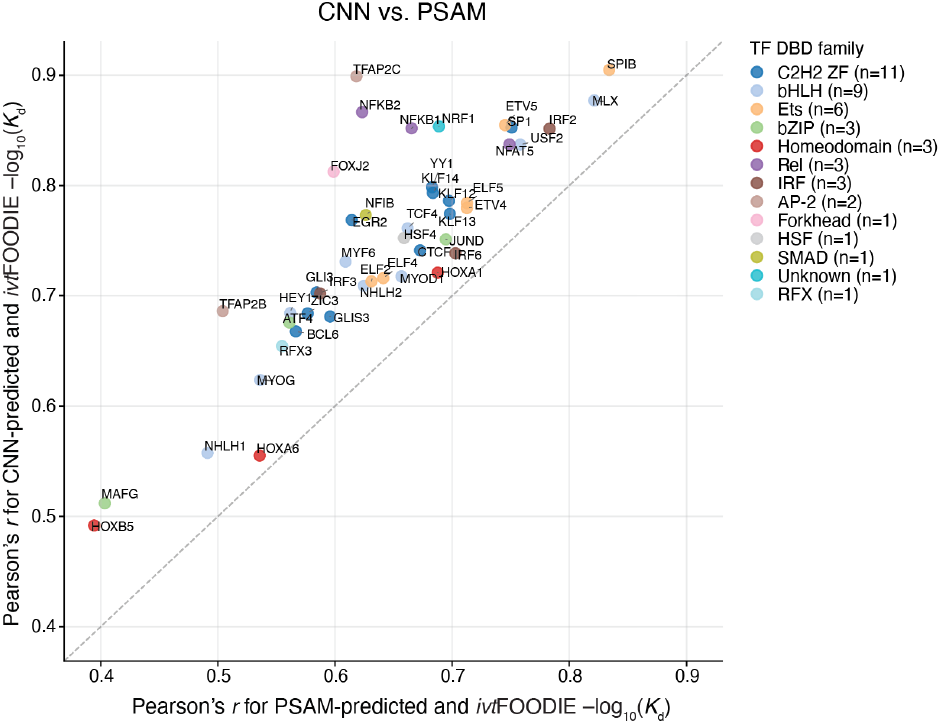
The effects of non-pairwise-additivity varies across TF families. Scatter plot comparing PSAM and CNN prediction performance for individual TFs. The x-axis shows Pearson’s *r* between experimentally measured affinities and PSAM-predicted affinities, whereas the y-axis shows Pearson’s *r* between experimentally measured affinities and CNN-predicted affinities. Each dot represents one TF, and dot color indicates TF family. Points above the diagonal indicate TFs for which the CNN outperforms the PSAM, reflecting improved prediction after relaxing the pairwise-additive assumption.

### Interdependency between Motif Positions Varies Substantially across TFs

To systematically map the spatial distribution of these non-pairwise-additive interactions, we used an *in-silico* double-mutant framework to compute the energetic epistasis (*ε*) between motif positions (Methods). Visualizing the signed pairwise dependency matrices revealed highly idiosyncratic architectural patterns (Figure 5a). The strongest long-range dependencies were predominantly observed in known obligate or facultative dimers (e.g., TFAP2C, NFKB1, JUND, and NRF1), directly reflecting energetic coupling between the two sub-monomer binding half-sites. The physical nature of this coupling varied dramatically: TFAP2C exhibited strong synergistic (positive) inter-monomer dependencies, whereas NFKB1 displayed pronounced antagonistic (negative) coupling. Conversely, TFs like SP1 and HEY1 lacked strong inter-site communication, with epistasis heavily concentrated at the core motif or strictly restricted to adjacent base pairs. Collectively, the matrices demonstrate that non-pairwise-additivity is a structurally-coupled, TF-specific thermodynamic signature.

**Figure 5.**
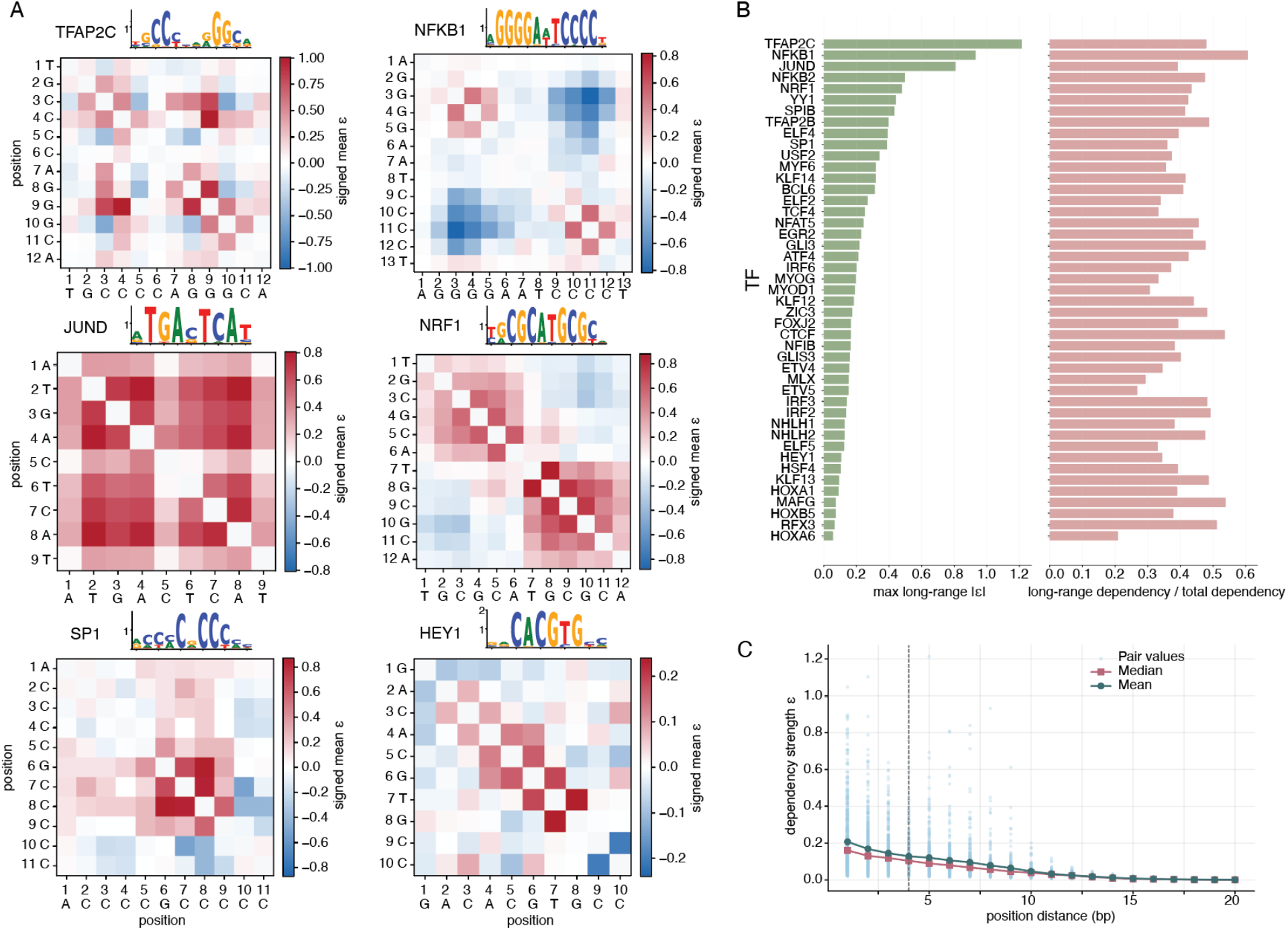
Interposition dependency patterns across TFs. (a) Representative signed mean pairwise dependency matrices for selected TFs. Heatmaps show the signed mean *ε* between motif positions, with red indicating positive dependency and blue indicating negative dependency. Sequence logos above each heatmap show the corresponding motif. TFAP2C, NFKB1, JUND, and NRF1 exhibit strong long-range dependency but with distinct patterns, whereas SP1 and HEY1 show more localized dependencies. (b) Long-range dependency across TFs. Long-range dependency was defined as dependency between positions separated by more than 4 bp. The left panel shows the maximum absolute long-range dependency strength for each TF. The right panel shows the fraction of total dependency contributed by long-range position pairs. TFs are ordered by maximum long-range dependency strength. (c) Dependency strength, *ε*, as a function of distance between motif positions. Each point represents the dependency strength for one pair of positions across TFs. The red line indicates the median dependency strength at each positional distance, and the blue line indicates the mean. Dependency strength generally decreases with increasing positional distance, but large non-pairwise-additive effects are still observed for some distant position pairs.

Focusing strictly on these distal dependencies, we quantified the maximum absolute epistasis and aggregate contribution to the total non-pairwise-additivity of long-range dependencies (separation >= 4 bp, Figure 5b). Strikingly, even for TFs lacking acute individual long-range interactions, distal coupling collectively accounted for up to 40% of the non-pairwise-additive energy landscape.

Broadly, as expected, the average dependency strength decayed rapidly with increasing distance between mutated positions, confirming that nearest-neighbor and short-range interactions generally dominate non-pairwise-additivity (Figure 5c). However, a distinct heavy-tail distribution emerged for distal pairs. While the mean long-range *ε* was modest (0–0.2), a critical subset of position pairs exhibited strong long-range epistasis with |*ε*| approaching 1.0. Given our – log_10_*K*_d_ scale, an |*ε*| of 1.0 translates to an order-of-magnitude shift in binding affinity, underscoring that distal co-mutations can exert large functionally significant synergistic or antagonistic energetic effects.

## DISCUSSION AND CONCLUSION

By integrating genome-wide thermodynamic *K*_d_ measurements with deep learning, we have systematically mapped the limits of the pairwise additive assumption in TF–DNA recognition. Our results provide definitive evidence that while pairwise additive frameworks (PWMs and PSAMs) capture essential binding motifs, they fundamentally fail to resolve a substantial portion of the affinity landscape. The superior performance of non-linear models, particularly CNNs, quantifies the energetic magnitude of these missing interactions, establishing non-pairwise-additivity as a defining feature of the TF–DNA binding energy landscape rather than a marginal correction.

Importantly, the extent of these non-pairwise-additive effects is intrinsically tied to the architectural logic of TF DBDs. Our data reveal a distinct functional spectrum: homodimer families like AP-2 (TFAP2C) and Rel (NFKB1) exhibit profound non-additivity, reflecting highly cooperative coupling between subunits, whereas HOX homeodomains adhere closely to the pairwise additive limit (Figure 4). As a striking manifestation of this complex energetic landscape, the widespread prevalence of long-range interposition dependencies observed in our *in-silico* screens provides a direct thermodynamic signature of distal energetic coupling (Figure 5). Such family-specific divergence implies that evolution has fine-tuned the exploitation of these energetic dependencies, allowing different TFs to achieve varying levels of regulatory precision through the selective use of non-pairwise-additive energetics.

Our framework directly relies on absolute thermodynamic *K*_d_, meaning the predicted epistasis represents true physical interaction energies rather than statistical correlations. This strict energetic mapping is what uniquely enables the resolution of subtle allosteric communications that typically evade traditional occupancy-based assays.

Several challenges remain. While our dataset spans 45 TFs, expanding to the entire human repertoire is essential for a comprehensive accounting of non-pairwise-additive DNA recognition. Bridging the gap between an *in vitro* biophysical baseline and the complexities of chromatin *in vivo*, including nucleosome positioning and co-factor competition, is a critical next step.

In summary, we established a rigorous quantitative deep learning framework that circumvented non-pairwise-additivity. This study provides quantitative models necessary to describe thermodynamics of TF-DNA interactions in genomic regulatory regions.

## Data Availability

Code for the models is available on GitHub (https://github.com/shenxiaoke/ivtFOODIE-CNN_dinucle_PSAM-train).

## ACKNOWLEDGMENTS

This project was financially supported by Changping Laboratory (Grants No. 2021C0402) and the Ministry of Science and Technology of China.

We thank Prof. Mingchen Chen (Changping Laboratory, Beijing, P. R. China.) and Prof. Jian Yan (Department of Biomedical Sciences, College of Biomedicine, City University of Hong Kong, Hong Kong SAR, China.) for helpful discussions.

## REFERENCES

(1) Lambert, S. A.; Jolma, A.; Campitelli, L. F.; Das, P. K.; Yin, Y.; Albu, M.; Chen, X.; Taipale, J.; Hughes, T. R.; Weirauch, M. T. The human transcription factors. Cell 2018, 172 (4), 650–665.

(2) Stormo, G. D. Modeling the specificity of protein-DNA interactions. Quantitative biology 2013, 1 (2), 115–130.

(3) Stormo, G. D. DNA binding sites: representation and discovery. Bioinformatics 2000, 16 (1), 16–23.

(4) Foat, B. C.; Morozov, A. V.; Bussemaker, H. J. Statistical mechanical modeling of genome-wide transcription factor occupancy data by MatrixREDUCE. Bioinformatics 2006, 22 (14), e141–e149.

(5) Schneider, T. D.; Stephens, R. M. Sequence logos: a new way to display consensus sequences. Nucleic acids research 1990, 18 (20), 6097–6100.

(6) Rauluseviciute, I.; Riudavets-Puig, R.; Blanc-Mathieu, R.; Castro-Mondragon, J. A.; Ferenc, K.; Kumar, V.; Lemma, R. B.; Lucas, J.; Chèneby, J.; Baranasic, D. JASPAR 2024: 20th anniversary of the open-access database of transcription factor binding profiles. Nucleic acids research 2024, 52 (D1), D174–D182.

(7) Vorontsov, I. E.; Eliseeva, I. A.; Zinkevich, A.; Nikonov, M.; Abramov, S.; Boytsov, A.; Kamenets, V.; Kasianova, A.; Kolmykov, S.; Yevshin, I. S. HOCOMOCO in 2024: a rebuild of the curated collection of binding models for human and mouse transcription factors. Nucleic Acids Research 2024, 52 (D1), D154–D163.

(8) Man, T.-K.; Stormo, G. D. Non-independence of Mnt repressor–operator interaction determined by a new quantitative multiple fluorescence relative affinity (QuMFRA) assay. Nucleic acids research 2001, 29 (12), 2471–2478.

(9) Bulyk, M. L.; Johnson, P. L.; Church, G. M. Nucleotides of transcription factor binding sites exert interdependent effects on the binding affinities of transcription factors. Nucleic acids research 2002, 30 (5), 1255–1261.

(10) Jolma, A.; Yan, J.; Whitington, T.; Toivonen, J.; Nitta, K. R.; Rastas, P.; Morgunova, E.; Enge, M.; Taipale, M.; Wei, G. DNA-binding specificities of human transcription factors. Cell 2013, 152 (1), 327–339.

(11) Fukushima, K. Neocognitron: A self-organizing neural network model for a mechanism of pattern recognition unaffected by shift in position. Biological cybernetics 1980, 36 (4), 193–202.

(12) LeCun, Y.; Boser, B.; Denker, J. S.; Henderson, D.; Howard, R. E.; Hubbard, W.; Jackel, L. D. Backpropagation applied to handwritten zip code recognition. Neural computation 1989, 1 (4), 541–551.

(13) Alipanahi, B.; Delong, A.; Weirauch, M. T.; Frey, B. J. Predicting the sequence specificities of DNA-and RNA-binding proteins by deep learning. Nature biotechnology 2015, 33 (8), 831–838.

(14) Zhou, J.; Troyanskaya, O. G. Predicting effects of noncoding variants with deep learning–based sequence model. Nature methods 2015, 12 (10), 931–934.

(15) Avsec, Ž.; Weilert, M.; Shrikumar, A.; Krueger, S.; Alexandari, A.; Dalal, K.; Fropf, R.; McAnany, C.; Gagneur, J.; Kundaje, A. Base-resolution models of transcription-factor binding reveal soft motif syntax. Nature genetics 2021, 53 (3), 354–366.

(16) Berger, M. F.; Philippakis, A. A.; Qureshi, A. M.; He, F. S.; Estep III, P. W.; Bulyk, M. L. Compact, universal DNA microarrays to comprehensively determine transcription-factor binding site specificities. Nature biotechnology 2006, 24 (11), 1429–1435.

(17) Jolma, A.; Kivioja, T.; Toivonen, J.; Cheng, L.; Wei, G.; Enge, M.; Taipale, M.; Vaquerizas, J. M.; Yan, J.; Sillanpää, M. J. Multiplexed massively parallel SELEX for characterization of human transcription factor binding specificities. Genome research 2010, 20 (6), 861–873.

(18) Barski, A.; Cuddapah, S.; Cui, K.; Roh, T.-Y.; Schones, D. E.; Wang, Z.; Wei, G.; Chepelev, I.; Zhao, K. High-resolution profiling of histone methylations in the human genome. Cell 2007, 129 (4), 823–837.

(19) Fordyce, P. M.; Gerber, D.; Tran, D.; Zheng, J.; Li, H.; DeRisi, J. L.; Quake, S. R. De novo identification and biophysical characterization of transcription-factor binding sites with microfluidic affinity analysis. Nature biotechnology 2010, 28 (9), 970–975.

(20) Nutiu, R.; Friedman, R. C.; Luo, S.; Khrebtukova, I.; Silva, D.; Li, R.; Zhang, L.; Schroth, G. P.; Burge, C. B. Direct measurement of DNA affinity landscapes on a high-throughput sequencing instrument. Nature biotechnology 2011, 29 (7), 659–664.

(21) Zhi Wang, D. W., Ke Shen, Junchen Luo, Xinyao Wang, Nan Wu, Yunzhi Lang, Xiangyu Wang, Jun Ren, Wenyang Dong, Lu Pan, Yitong Lyu, Gang Li, Dubai Li, Chen Xie, Zhen Zhang, Shijun Yu, Liuying Shan, Nannan Zhang, Jian Yan, Mingchen Chen, Xiaoliang Sunney Xie. Prediction of Transcription Factor DNA Binding Affinity with High-Throughput Kd Measurements and Deep Learning. bioRxiv 2026.

(22) Buenrostro, J. D.; Giresi, P. G.; Zaba, L. C.; Chang, H. Y.; Greenleaf, W. J. Transposition of native chromatin for fast and sensitive epigenomic profiling of open chromatin, DNA-binding proteins and nucleosome position. Nature methods 2013, 10 (12), 1213–1218.

(23) Hill, A. V. The possible effects of the aggregation of the molecules of hemoglobin on its dissociation curves. j. physiol. 1910, 40, iv-vii.

(24) Bailey, T. L.; Johnson, J.; Grant, C. E.; Noble, W. S. The MEME suite. Nucleic acids research 2015, 43 (W1), W39–W49.

